# Database Chemistry for Genomics-Based Safety and Quality Evaluation of Biologics

**DOI:** 10.1101/2025.02.14.638017

**Authors:** Norichika Ogata, Tomoko Matsuda, Aoi Hosaka, Akihisa Shina, Noriko Hashiba, Kazuhisa Uchida, Yoshinori Kawabe, Masamichi Kamihira, Takanori Yamaoka, Hiromu Kurita, Noriko Yamano-Adachi, Takeshi Omasa

**Affiliations:** Nihon BioData Corporation, Kawasaki, Kanagawa, 213-0012, Japan; Graduate School of Engineering, Osaka University, Suita, Osaka, 565-0871, Japan; Manufacturing Technology Association of Biologics, Chuo-ku, Tokyo, 104-0033, Japan; Faculty of Engineering, Kyushu University, Fukuoka, Fukuoka, 819-0395, Japan; Graduate School of Science, Technology and Innovation, Kobe University, Kobe, Hyogo, 650-0047, Japan

**Keywords:** CHO cells, Viral Safety, Host Cell Proteins, Genome, Single Cell RNA-seq, Database, SWATH-MS

## Abstract

Genomics-based safety and quality evaluation studies are advancing the bioindustry by enhancing various aspects, including viral safety, host cell protein (HCP) control, product heterogeneity control, cellular heterogeneity control, and process reproducibility. High-throughput instruments and genome-scale databases are essential in genomics, with the reference genome sequence being the most critical database. The completeness and accuracy of these genome sequences depend on DNA quality, sequencing instruments, read coverage, and assembly strategies. Significant efforts are being made to perfect genome assembly and continuously improve it. However, the quantitative impact of reference genome sequence accuracy on the safety of biologics is not yet fully understood. In this study, we compared and benchmarked six Chinese hamster genomes, including four newly sequenced genomes derived from Chinese hamster cell lines, from an industrial perspective. We also developed database assembly techniques to enhance the safety of biologics. We recommend using two or more independent reference genomes for viral safety studies. For HCP control, we suggest using protein sequences in which trypsin degradation peptides that overlap with high-risk proteins should be masked and unified. Additionally, we can predict microenvironments using single-cell transcriptome data. In bioengineering processes, any nucleotide samples have potential commercial benefits.

## I. Introduction

Genomics, which relies on genome-scale databases and high-throughput instruments, is at the forefront of scientific research and industrial advancements. Recently, our study, which is the first report on mutant strains of a pandemic virus, SARS-CoV-2, demonstrated the scientifically, medically, and economically significant application of genomics: tracing infection routes (Matsuda et al., 2020). Genomics should become indispensable in the future production of biologics, enhancing viral safety studies, host cell protein control, product heterogeneity control, cellular heterogeneity control, and process reproducibility. Genome sequences are the most fundamental databases in genomics, and the accuracy of these assemblies has significantly improved in the past decade (Ogata, 2021). There are numerous assemblies of the same organisms, including both published and private genomes (Birzele et al., 2010; Rupp et al., 2018; Xu et al., 2011), allowing us to select assemblies based on the specific purpose of our analyses. However, it is unclear how to select the appropriate databases and to justify these choices. The most accurate genome may not necessarily be the best option, as several studies have evaluated genome assemblies (Manchanda et al., 2020; Mikheenko et al., 2023; Wang & Wang, 2023). In this study, we sequenced and assembled the genomes of four Chinese hamster-derived cell lines. We analyzed laboratory-scale cellular datasets using these genomes and the databases derived from them. The scope of this study encompasses five major topics: viral safety studies, host cell protein control, product heterogeneity control, cellular heterogeneity control, and process reproducibility studies. We introduce these topics, focusing on their purposes, methods, and the risks of false negatives and positives.

Historically, viral safety studies (Barone et al., 2020; Merten, 2002), have involved in vivo and in vitro infectious examinations, as well as observations using electron microscopes. The international guideline ICH Q5A (ICH, 1998a, 1998b), is recently revised; it now acknowledges the applicability of Next Generation Sequencing (NGS) technology for broad and specific virus detection. In the past decade, detecting infectious elements in animal tissues using NGS has become a popular technique in genomics (Kostic et al., 2011). In these studies, infectious elements are identified as sequences that cannot be mapped to the host genome. However, if the host genome is incomplete, some host cell-derived virus-like sequences could mistakenly be detected as infectious elements due to the presence of many virus-like sequences in the genomes of multicellular organisms. In this study, we conducted viral safety analyses using Chinese hamster cells spiked with viruses and several public and in-house Chinese hamster genomes. We also addressed two major issues. The first issue is the undeniable possibility of viral contamination in the cells used as a source for the reference genome (Baker, 2012; Yoshida et al., 2017). Studies have reported that 3.3% to 31.7% of cell lines are contaminated with viruses (Bolin et al., 1994; Uphoff et al., 2019). If the cells used to produce the reference genome are contaminated, viral sequences might be incorrectly classified as part of the host cell genome, leading to a risk of false-negative results. The second issue is read contamination between samples. Even with meticulous biological sample handling and fully robotic automated sample preparations, there remains a risk of imperfect demultiplexing of reads in NGS flow cells (Nieuwenhuis et al., 2020). Very careful bio sample manipulations and totally robotical automated sample preparations cannot certificate imperfect demultiplexing of reads in flow cells of NGS. To mitigate this issue, we attempted to eliminate contamination by using single-use NGS without multiplexing (“Nanopore Sequencing,” 2024).

In the host cell protein (HCP) control study, immunoassays like enzyme-linked immuno-sorbent assay (ELISA) were performed historically using polyclonal anti-host cell antibodies (Graf et al., 2021). In recent years, lists of critically harmful host cell proteins have been established (Jones et al., 2021) and it has become possible to quantify each protein with reference to risk knowledge using LC-MS/MS technologies (Carvalho et al., 2024; Oh et al., 2024). In this study, we conducted an HCP control study using Chinese hamster cells and protein databases derived from several public and in-house Chinese hamster genomes. In general, comparative proteomic analyses, the most important data points are the differences between biological samples of the same proteins, where all quantified subjects hold equal biological value. However, in the context of HCP control, the value of quantified subjects varies significantly, as the false-negative detection of high-risk HCPs must be strictly avoided. In the proteomic measurement process, an ion library database is assembled from protein sequences derived from genomes and Data-Dependent Acquisition (DDA) using LC-MS/MS. Peptides with the same molecular weight and time of flight cannot be distinguished in MS/MS, so duplicated protein sequences are typically excluded from the ion library. General protocols often overlook partial protein sequences of high-risk HCPs that are shared with other proteins. To reduce the risk of false negatives for high-risk HCPs, we aligned the protein sequences and unified the duplicated protein sequences. We then examined these database assemblies using cells derived from Chinese hamsters.

In the products heterogeneity control study (Cartwright et al., 2018), we can find mutations in product proteins from single cell RNA-seq (scRNA-seq) data of subjects producing cells. Generally, humanized antibody sequences are not included in the host genome and no special technique is required. In this study, we searched for humanized antibody-like sequences in public and in house Chinese hamster genomes.

In the cellular heterogeneity control study, there are two major purposes; The first purpose is to observe the diversity of a culture and the second is to prove the monoclonality of cells (Borsi et al., 2023; Ogata et al., 2021; Tzani et al., 2021). The first purpose is very similar to other scRNA-seq projects; We can draw attractive colorful plots of results of the principal component analysis, the t-SNE or the UMAP. In these projects, we are attempting to understand cellular diversity. We demonstrated scRNA-seq and diversity visualization in three Chinese hamster derived cell lines using publicly available and in house genomes. The second purpose was proposed in our previous study; mitochondrial single nucleotide variations work as the clonal marker. We performed several single cell sequencing protocols and surveyed. For the process reproducibility studies, RNA-seq would be the most appropriate while there are several monitoring methods available including Raman spectroscopy (Tanemura et al., 2023; Yan et al., 2024). PCA and similar methods would be useful for checking the process reproducibility. Here, we developed machine learning models between concentrations of chemical components and RNA-seq derived quantitative genome expression data using publicly available and in house genomes. We also performed single cell scale micro environmental estimation (Ogata et al., 2018) using single cell RNA-seq data.

## II. Materials AND METHODS

### Cell Culture

We cultured 4 cell lines, the CHO-K1 cells, CHO-S cells, CHL-YN cells (Yamano-Adachi et al., 2020) and CHK-Q cells (Kawabe & Kamihira, 2022). The CHO-K1 cells were obtained from ATCC and cultured in 20 mL of EX-CELL CD CHO Fusion (14365C-1000ML; Merck, Darmstadt, Germany) containing 600µ L of 200 mmol/L L-Glutamine Solution (x100) (073-05391, Fujifilm Wako Pure Chemical, Osaka, Japan) in 125 mL flasks. The flasks were shaken (φ25 mm, 120 rpm). The CHO-S cells were obtained from ATCC and cultured in 20 mL of FreeStyle CHO Expression Medium (14365C-1000ML; Gibco, Grand Island, NY, USA) containing 600 µL of 200mmol/L L-Glutamine Solution (x100) (073-05391, Fujifilm Wako Pure Chemical, Osaka, Japan) in 125 mL flasks. The flask was shaken (φ25 mm, 140 rpm). The CHL-YN cells which was established in our laboratory in the Osaka university were cultured in 20 mL of EX-CELL CD CHO Fusion (14365C-1000ML; Merck, Darmstadt, Germany) containing 600 µL of 200 mmol/L L-Glutamine Solution (x100) (073-05391, Fujifilm Wako Pure Chemical, Osaka, Japan) in 125 mL flasks. The flasks were shaken (φ25 mm, 120 rpm) at 37 °C in a humidified 5% CO^2^ atmosphere. The CHK-Q cells which was established in our laboratory in the Kyushu university were cultured in DMEM/F-12 (DF) medium (042_30555; Fujifilm Wako Pure Chemical, Osaka, Japan) supplemented with 10% fetal bovine serum (FBS) (Biowest, Nuaill, France) in 100-mm cell culture dishes coated with collagen type I (4020_010; AGC Techno Glass, Shizuoka, Japan). The CHL-YN cells producing an IgG_1_ were also perfusionally cultured in EX-CELL Advanced HD Perfusion Medium (MERCK) containing L-glutamine (final concentration: 6 mM; Wako, 073-053911) and Puromycin (final concentration, 5μg/mL; Invivogen) at 37 °C, 80% humidity, 5% CO2, and 120 rpm (φ25 mm) in a shaker before inoculation in the bioreactor. Perfusion process runs were performed in 2 L BCP-07NP3 bioreactor (ABLE Biott, Tokyo, Japan) with XCELL™ ATF2 system. The diameter of the impeller is 55 mm. The hollow fiber filter (HF) was Filter 0.2-µm polyethersulfone (Repligen). The bioreactor was incubated at 37 °C, pH 7.2, 120 rpm, and DO was maintained at 3.43 ppm. The perfusion was started at 1.2 RV/day (0.8 L/min) on day 2.5 using EX-CELL Advanced HD Perfusion Medium (MERCK) with 15 mM L-glutamine, 5 μg/mL Puromycin and 0.1% anti-foaming agents. The exchange rate was changed to 1.2 L/min (2.0 RV/day) on day 3. The bleeding started on day 3 and the viable cell density was kept at 2.0 × 10^7^ cell/mL. The glutamine and glucose concentration were controlled at 2 mM and 3 g/L, respectively.

### Genome Sequencing and Assemble

Methods were previously described (Ogata, 2021). Cellular genomic DNA samples were extracted using the Qiagen Blood & Cell Culture DNA Maxi Kit (Qiagen, Gaithersburg, MD). The concentrations of cells and viabilities of the CHO-K1 cells, the CHO-S cells, the CHL-YN cells and the CHK-Q cells were 56.4 × 10^5^ cells/mL (98.6 %), 1.35 × 10^6^ cells/mL (95.0 %), 1.14 × 10^7^ cells/mL (96.3 %) and 1.35 × 10^6^ cells/mL (95.0 %), respectively. We used 1.0 × 10^8^ cells in all the cell lines. The DNA quality and quantity were measured using the TapeStation. All DNA samples were prepared for sequencing using Sequel II Binding Kit 2.0 (Pacific Biosciences) and Sequel II DNA Internal Control Kit 1.0 (Pacific Bio-sciences). Sequencing was done on the Sequel II System (Pacific Biosciences) using Sequel II Sequencing Kit 2.0 (Pacific Biosciences) and Sequel SMRT Cell Oil (Pacific Biosciences). Data processing was performed using SMRT Link v8.0.0 (Pacific Biosciences). Library preparation, sequencing and base calling were performed at Macrogen Japan Corp. (Tokyo, Japan). The Hi-Fi reads were assembled using the Hifiasm (v 1. 6) program. The super-computing resource, SHIROKANE, was provided by Human Genome Center, the Institute of Medical Science, the University of Tokyo.

### Epigenomic analyses

Illumina bisulfite sequencing libraries were prepped using ACCEL-NGS® METHYL-SEQ (Swift Biosciences, Inc., Michigan, US) and sequenced using Novaseq 6000 (Illumina, Inc., SD, US). The reads were 150bp paired. The reads were processed using Bismark v0.24.2.

### Transcriptome sequencing (RNA-seq)

Methods were previously described (Matsuda, 2021). The QIA shredder and the RNeasy were used. Illumina libraries were prepped and sequenced using Novaseq 6000. For a RNA-seq using a single use NGS, double-stranded cDNA was synthesized using NEBNext Ultra™ II Directional RNA Library Prep Kit for Illumina (NEB) following manufacturer’s protocol with several modifications. Firstly, 500 to 1,000 µg of input RNA is recommended; however, for this study, 8000 ng was used. Accordingly, the volume of the double-stranded cDNA synthesis kit was increased eightfold. The reaction was conducted in eight separate tubes. To amplify the double-stranded cDNA, a 15-cycle PCR reaction was performed. Eluted DNA was quantified using Qubit 3.0 (Thermo Fisher Scientific).

The DNA was then used to prepare a Nanopore-sequencing library using DNA ligation Kit (SQK-LSK109: Oxford Nanopore Technologies) with several modificaitons. Firstly, 2,500 ng of DNA was used for the library preparation. Secondly, during DNA Repair and end-prep step, the reaction time was extended from 20°C for 5 minutes, then 65°C for 5 minutes to 20°C for 30 minutes, then 65°C for 30 minutes. Thirdly, during Adapter Ligation and Clean-up step, the DNA-bound AMpure XP beads (BECKMAN COULTER) was washed 250 μl of Short Fragment Buffer, instead of Long Fragment Buffer. All the eluted library was sequenced and basecalled using FLO-MIN106D Flow Cell (R9.4.1) with MinION Mk1C.

### Single cell transcriptome sequencing (scRNA-seq)

The scRNA-seq was performed using the CHO-K1 cells (C1), the CHO-S cells (C1) and the CHL-YN cells (C1, 10X Chromium and BD Rhapsody). In the process of Fluidigm C1, single cell loading and capture were performed following the Fluidigm protocol (PN 100-7168). Briefly, 40 μL of C1 Suspension Reagent was added to a 60 μL suspension of 400,000 cells in the PBS(-) with 5 mg/mL bovine serum albumin (BSA) filtered with 35 µm Cell-Strainer. Pipet 6 μL of the cell mix into the cell inlet of C1 IFC for mRNA Seq (10–17 μm, PN 100-5760). Place the IFC into the C1. Run the SMART-Seq v4 Cell Load Rev B script. Then, running lysis, reverse transcription, and PCR were performed on the C1 system. Library preparation was performed with a Nextera XT DNA Library Preparation Kit (Illumina, FC-131-1096) following the Fluidigm protocol. The finished cDNA libraries were quantified by using the Agilent 2200 TapeStation and sequenced on an Illumina NextSeq 500 platform with 75-bp single-end reads.

The scRNA-seq using the 10X Chromium system 10X Genomics) was performed by the Research Institute for Microbial Diseases, Osaka University. CHL-YN cells used for the Fluidigm C1 and BD Rhapsody experiments were loaded onto a ChromiumX. Suspended CHL-YN cells were filtered with 35 µm Cell-Strainer in the PBS(-) with 0.04% BSA. Cells were processed following Chromium Single Cell 3’ Reagent Kits protocol. Approximately 16,500 live cells per sample were loaded onto the Chromium Controller to generate 10,000 single-cell gel-bead emulsions for library preparation and sequencing. Libraries were sequenced on a DNBSEQ-G400RS (MGI).

The scRNA-seq using the BD Rhapsody system was performed by the Research Institute for Microbial Diseases, Osaka University. CHL-YN cells used for the Fluidigm C1 and 10X Chromium experiments were loaded in parallel onto a BD Rhapsody cartridge with a target capture of 10,000 cells. Suspended CHL-YN cells were filtered with 35 µm Cell-Strainer in the sample buffer. Cells were processed following the manufacturer’s guidelines of the BD Rhapsody Single-Cell Analysis System (BD Rhapsody). Libraries were sequenced on a DNBSEQ-G400RS (MGI).

### Databases assemble

We performed annotation of genomes; genes were found in the genomes and trans elements were found. The genome annotation of genes was performed using RNA-seq. The reads were aligned to the assembled genomes using hisat2 (Kim et al., 2019). The resulting BAM files were utilized to predict transcription units by StringTie2 (Kovaka et al., 2019). The resulting GTF files were then used to predict protein-coding regions by TransDecoder (v5.7.0). Genomes and annotations of the CHO-K1 cells were (CriGri-1.0, NCBI RefSeq: GCF_000223135.1 (Xu et al., 2011) and CriGri-PICRH-1.0, NCBI RefSeq: GCF_003668045.3 (Rupp et al., 2018)) obtained from the NCBI nucleotide database. The annotations on the PIRCH-1.0 were used to predict transcription units and homology search with LiftOff (Shumate & Salzberg, 2021). The RNA-seq based ab initio annotations were combined with those of based homology search with MAKER v3.01.03 (Campbell et al., 2014). The genomes and gene annotations were combined to transcripts sequences and protein sequences using gffreads. The annotation of trans elements was performed using RepeatMasker v4.1.1 (Tarailo-Graovac & Chen, 2009). The protein sequences obtained from the CriGri-PIRCH-1.0 and their trypsin degradation peptide shared with the 86 high risk proteins listed previously (Jones et al., 2021) were masked and unified.

### LC-MS/MS measurement

For the supernatant of CHL-YN cells, the medium containing the suspended cells was centrifuged for 10 min at 2,000 xg and the supernatant was concentrated by acetone precipitation. Protein concentration of the pellets that resuspended in PBS (-) was determined using the Pierce™ Modified Lowry Protein Assay Kit (Thermo Fisher Scientific) according to the manufacturer’s instructions. The pellets including 200 µg protein were resuspended in 100 µL of 6.4 M guanidine hydrochloride in 50 mM Tris hydrochloride (pH 7.9) and vortexed. Add 2 µL of 500mM dithiothreitol (DTT, 500mM) and incubate for 30 min at room temperature under dark conditions. For alkylation, add 4 µL of 500mM iodoacetamide (IAA) and incubate for 20 min at room temperature under dark conditions. Add 2 µL of 500mM DTT and incubate for 30 min at room temperature under dark conditions. Zeba™ Spin Desalting Columns 7K MWCO (Thermo Fisher Scientific) were used for the buffer exchange to Sodium bicarbonate (NaHCO_3_) and desalted according to the manufacturer’s instructions. Add 20uL trypsin solution (1mg/mL) to the sample and incubate for 18h at 37°C under dark conditions. After digestion, 10% FA was added to the sample to terminate the reaction and the reaction solution was filtered through a 0.22-μm syringe filter for LC-MS analysis.

For LC–MS/MS analysis, an estimated 200 µg of protein samples from each fraction were analyzed on a ZenoTOF 7600 mass spectrometer (SCIEX) coupled to a TurboIonSpray Ion Source (SCIEX) and interfaced with a Nexera X3 UHPLC System (Shimazu). The peptide samples were separated at 50 °C on an ACQUITY UPLC CSH C18 column (2.1 × 50 mm, particle size of 1.7 µm, and pore size of 130 Å). The mobile phase consisted of 0.1% formic acid in water (buffer A) and 0.1% formic acid in acetonitrile (buffer B). The column was maintained at 50°C, with a flow rate of 0.3 mL/min. The chromatographic gradient lasted 120 min, incorporating a linear increase from 2% to 35% B over 93 minutes, followed by a column wash (8 minutes at 90% B) and re-equilibration (13 minutes at 2% B). For data-dependent acquisition (DDA), MS/MS spectra were acquired for the 20 most intense precursor ions detected in each 250 ms MS survey scan, covering a mass range of 400–1250 m/z and considering precursor charge states ranging from 2 to 5. To minimize repeated selection of the same precursor, a dynamic exclusion window of 12 seconds was applied following a single detection.For sequential window acquisition of all theoretical mass spectra (SWATH) data-independent acquisition (DIA), 33 variable isolation windows (in Da) were employed over the 400–1250 m/z range.

### Data analyses for the viral safety study

The RNA-seq reads from four cell line samples were mapped to six reference genomes to remove sequences derived from host cell genomes using hisat2 v2.2.0. Then, we mapped the descents to the clustered version of the FDA reference viral genomes database (C-RVDB v24.1) and counted and normalized using the bowtie2 v2.4.1. and the mapping results were validated using the blastn v2.10.0 with-task blastn-short option. For virus contaminated model samples which were spiked in Moloney Murine Leukemia Virus (M-MuLV) at five concentrations, reads which were aligned to the MuLV genomes (K02728.1, NC_001501.1, NC_001502.1 and AH002383.2) were counted and summed. The statistical significance of the effect of reference genomes was tested by lm function in R v4.1.2 (R Core Team, 2024). The usage was summary(lm(objectiveValue∼referencialGenome)).

### Data analyses for the host cell proteins control study

We generated Chinese hamster (CH) spectral libraries and performed SWATH-MS data analyses. The spectral library was constructed using the workflow established by SCIEX. Briefly, the raw DDA data files were processed using the ProteinPilot software v5.0.2.0 against a protein sequence database. The database was the latest release of CH RefSeq assembly (CHO - K1, GCF _ 003668045.1), Cri Gri - 1.0 (CHO - K1, GCF_000223135.1), BAABOA000000000.1 (CHO-K1), BAABNZ 000000000.1 (CH O - S) a n d BAABNY000000000.1 (CHL-YN). The search effort setting was Rapid ID with carbamidomethyl (C) as a fixed modification, and oxidation (M), deamidation (NQ) and pyroglutamic acid conversion (EQ) as variable modifications. Each of the DDA raw data files was searched in ProteinPilot and the result files were used as the input libraries for spectral library generation. For the construction of the spectral library, each input library was filtered by 1% FDR and 99% confidence threshold to remove low-confidence peptide identifications.

The spectral libraries were constructed for each cell in the ProteinPilot software. The SWATH-MS data analysis was performed in the PeakView v2.2 on a local PC. In the local SWATH-MS processing workflow, the spectral library and SWATH-MS acquisition data were loaded into SWATH Processing microApp in PeakView v2.2. The peak groups were extracted with a 99% peptide confidence threshold and a 1% peptide FDR cutoff. The RT extraction window and the fragment ion mass tolerance were set to 5 min and 75 ppm, respectively. The statistical significance of the effect of reference genomes was tested by lm function in R v4.1.2. The us age was summary(1m (objective Value ∼ referencialGenome)). The statistical significance of the effect of the database modification was tested by fisher.test function in R v4.1.2.

### Data analyses for the product heterogeneity control study

The homology search between CH genomes and human IgG was performed using the BLAST v2.10.0.

### Data analyses for the cellular heterogeneity control study

The scRNA-seq read from three cell lines (the CHO-K1, CHO-S and CHL-YN cells) obtained from the C1 were mapped to the six reference genomes using the STAR v2.7.11b (Dobin et al., 2013) and counted using htseq-count v2.0.5. The PCA analyses were performed using the prcomp function on the R v4.1.2. The scRNA-seq reads from the CHL-YN cells obtained from the 10x Chromium were processed using cellranger v7.1.0. The scRNA-seq reads from the CHL-YN cells obtained from the Raphsody were processed using cwl-runner v.3.1.20230906142556 in the bdgenomics_rhapsody:2.0 on the singularity v.3.6.4.

### Data analyses for the process reproducibility studies

The RNA-seq obtained from cultured cells were checked using FastQC. The sequences were mapped to six genomes using salmon v1.10.2 (Patro et al., 2017) and translated to quantitative transcriptome data (TPM). These TPM data were analyzed with PCA. The quantitative transcriptome data were randomly divided into two datasets in a 9:1 ratio using random numbers for machine learning analyses. The prior data sets were used for training and the others were used for evaluating the prediction models. In this analysis, the explanatory variable was quantitative transcriptome and the object variable was mean cell size (µm), culture time (days), glucose concentration (g/L), pCO_2_ (mmHg), pH and pO_2_ (mmHg). We also predicted those object variables in micro environments from scRNA-seq data using the models.

### Ultrastructures observations

Methods were previously described (Ogata & Iwabuchi, 2017). Cells in media at 37 °C were fixed using the same volume 0.1 M phosphate buffer (pH 7.4) containing 4% paraformaldehyde and 4% glutaraldehyde at 37 °C. The fixed cells were kept at 4 °C. The cells were fixed overnight in a 0.1 M phosphate buffer (pH 7.4) containing 2% glutaraldehyde at 4 °C. The cells were washed three times for 30 minutes using 0.1 M phosphate buffer. The cells were fixed for 90 minutes in a 0.1 M phosphate buffer (pH 7.4) containing 2% osmium tetroxide at 4 °C. The cells were dehydrated in 50 % ethanol at 4 °C for 10 minutes, 70 % ethanol at 4 °C for 10 minutes, 90 % ethanol at room temperature for 10 minutes and absolute ethanol at room temperature for 10 minutes three times. The cells were plastinated in propylene oxide at room temperature for 30 minutes twice, a 7:3 solution of propylene oxide and Quetol-812 (Nisshin EM Co., Tokyo, Japan) at room temperature for 1 hour. The propylene oxide was volatilised at room temperature overnight. The Quetol-812 containing the cells was fixed at 60 °C for 48 hours. Ultrathin sections (70 nm) were cut using Ultracut UCT (Leica, Vienna, Austria) using a diamond knife. The sections were scooped up on Cu grids. The sections were dyed in 2 % uranyl acetate at room temperature for 15 minutes and in a lead stain solution (Sigma-Aldrich Co.) at room temperature for 3 minutes. Ultrastructures of the cells were observed using a transmission electron microscopy at 100kV, JEM-1400 Plus (JEOL Ltd., Tokyo, Japan). The images were captured in 3296 × 2472 pixels (9.285 µm × 6.963 µm) using a CCD camera, EM14830RUBY2 (JEOL Ltd.).

## III. Results

### Genomes sequencing, assembly and annotation

Genomic DNA samples were extracted from the four Chinese hamster derived cell lines, the CHO-K1, CHO-S, CHL-YN (Yamano-Adachi et al., 2020) and CHK-Q (Kawabe & Kamihira, 2022); The DNA was dissolved in 1 ml of TE buffer with the concentration of 381 ng/ul, 480 ng/ul, 370 ng/ul,and 224 ng/ul, respectively. The integrity of the DNA samples was confirmed using Pulse-Field Gel Electrophoresis (0.8% Agarose Gel, Wave-from Type: 5-80K, Run Time: 17 hrs). HiFi long reads of the genomic DNA from the CHO-K1, CHO-S, CHL-YN, and CHK-Q cell lines were obtained using the PacBio Sequel IIe system in CCS mode. The number of reads and N50 values for CHO-K1, CHO-S, CHL-YN, and CHK-Q were 8,072,185 (20,352 bp), 9,155,380 (16,697 bp), 8,539,963 (18,387 bp), and 8,548,111 (18,759 bp), respectively. The total nucleotides obtained from the CHO-K1, CHO-S, CHL-YN and CHK-Q cells were 161.4, 153.5, 156.1 and 157.9 Gb, respectively. Reads were assembled using Hifiasm v.0.16.1-r375. The coverages and genome sizes of the CHO-K1, CHO-S, CHL-YN and CHK-Q cells estimated by Hifiasm were x63 (2.6 Gb), x58 (2.6 Gb), x57 (2.7 Gb) and x58 (2.7 Gb), respectively. The assembled genome size and contig N50 of the CHO-K1, CHO-S, CHL-YN and CHK-Q cells were 2.94 Gb (41.0 Mb), 2.78 Gb (37.8 Mb), 2.69 Gb (83.0 Mb) and 2.64 Gb (112.5 Mb), respectively. The contig number of the CHOK1, CHO-S, CHL-YN and CHK-Q cells were 3133, 2540, 600 and 457, respectively. The genome sequences are available from GenBank; The accession numbers of the genomes are BAABOA000000000.1 (CHO-K1), BAABNZ000000000.1 (CHO - S), BAABNY000000000.1 (CHL-YN) and BAABNX000000000.1 (CHK-Q). We also obtained two public genomes of the Chinese hamster derived cell lines, CriGri-1.0 (CHO-K1, GCF_000223135.1) and CriGri-PICRH-1.0 (CHO-K1, GCF_003668045.3). The obtained reads from bisulfite sequencing of the CHO-K1, CHO-S, CHL-YN and CHK-Q cells were 434,400,987, 437,541,571, 445,183,944 and 435,823,715, respectively. The methylation rates of CpG islands were highest in the following order: the CHO-K1, CHOS, CHL-YN and CHK-Q in whole genome bisulfite sequencing analyses (Supplemental Figure 1).

**Supplemental Figure 1.**
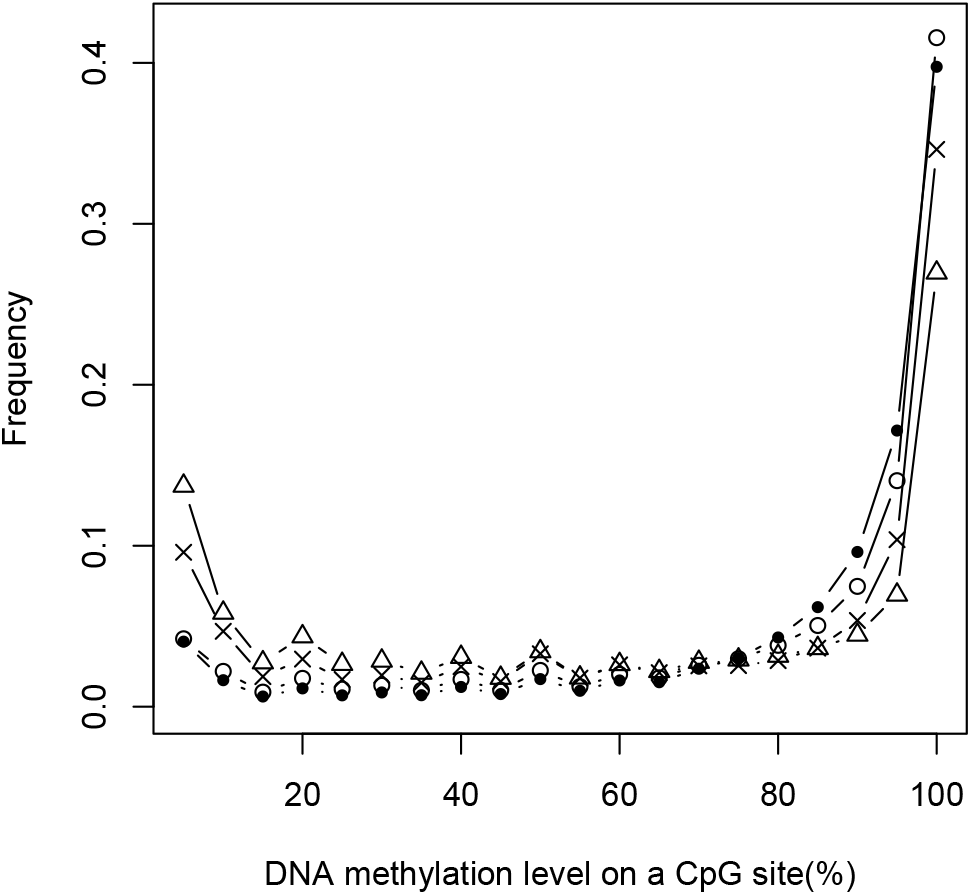
Frequency of DNA methylation level on CpG sites in four cell lines. Open circles, closed circles, crosses and open triangles indicate the CHO-K1, CHO-S, CHL-YN and CHK-Q cells, respectively.

We annotated the newly obtained genomes in three methods, ab initio annotation using RNA-seq data, homology search using the CriGri-PICRH-1.0 annotation and combinations of the two annotations (ab initio and homology searching). We choose annotations depending on the purpose. In addition, we annotated transposable elements, including virus-like sequences, and compared their expression across different cell lines. The variation in transposable element expression between cell lines was smaller than the variation observed in gene expression (Supplemental Figure 2).

### Viral Safety Studies

The number of normalized reads mapped to the model virus genome (M-MuLV) correlated with the spiked-in concentration, regardless of which genome was used to filter out host-derived reads (Figure 1a). The percentage of reads that mapped neither to the host genome nor to the viral database, relative to the total reads used in the analysis, was highest when GriGri-1.0 was used (mean: 9.89%, median: 9.22%, min: 7.62%, max: 15.10%), followed by CriGri-PICRH-1.0 (mean: 4.82%, median: 4.80%, min: 2.92%, max: 6.54%), and was extremely low when our newly determined four genomes were used (mean: 0.98%, median: 0.98%, min: 0.75%, max: 1.35%) (Figure 1b). Using the newly sequenced, highly complete genome as a reference genome reduced (p = 0.000012, regression analysis) the number of unidentified reads that potentially contain unknown viruses or other contaminants.

**Figure 1.**
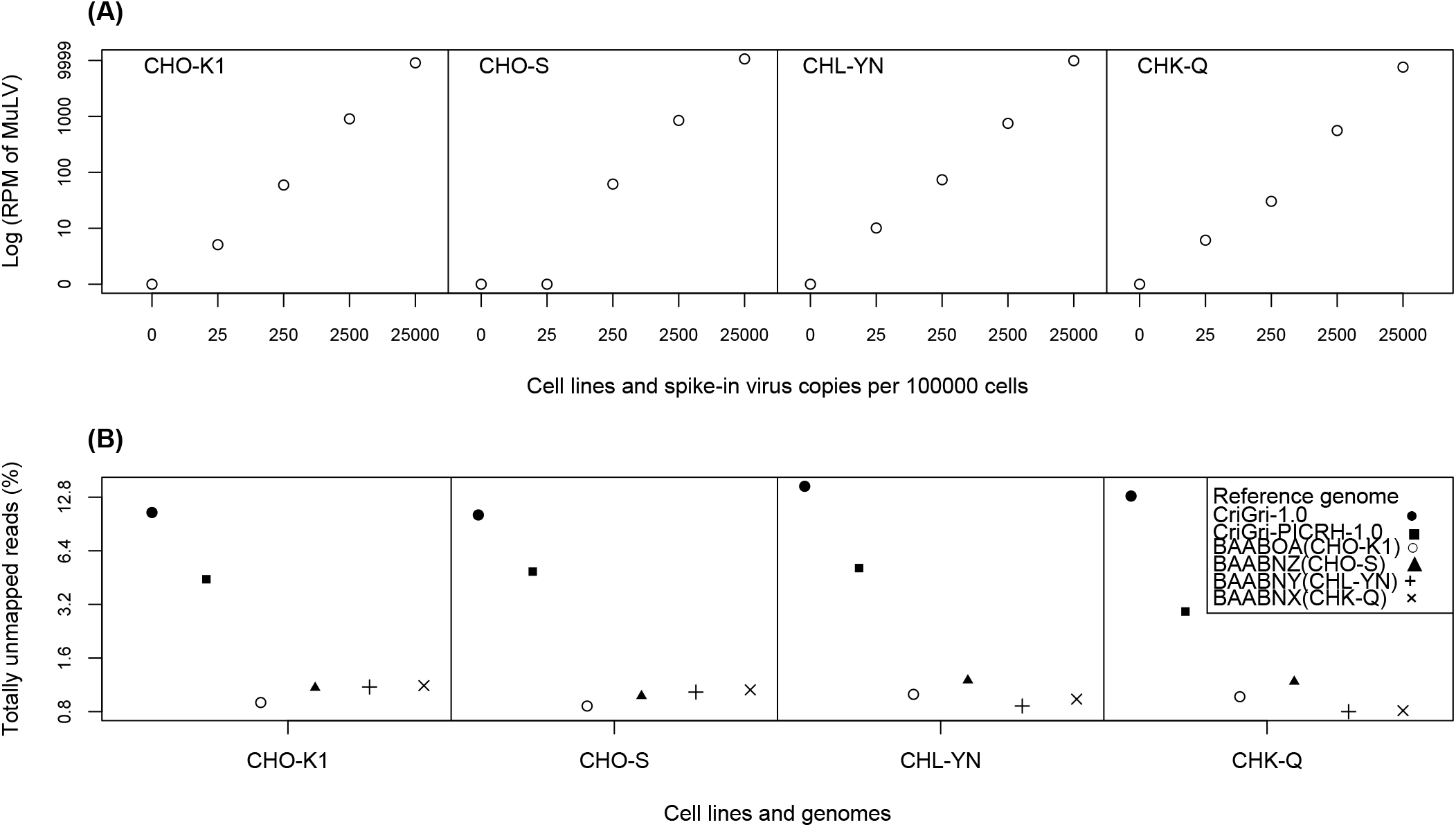
Viral safety studies using multiple genomes. A) Quantification of spiked viral sequences across different cell lines. (B) Comparison of unmapped read rates among various reference genomes across different cell lines.

**Supplemental Figure 2.**
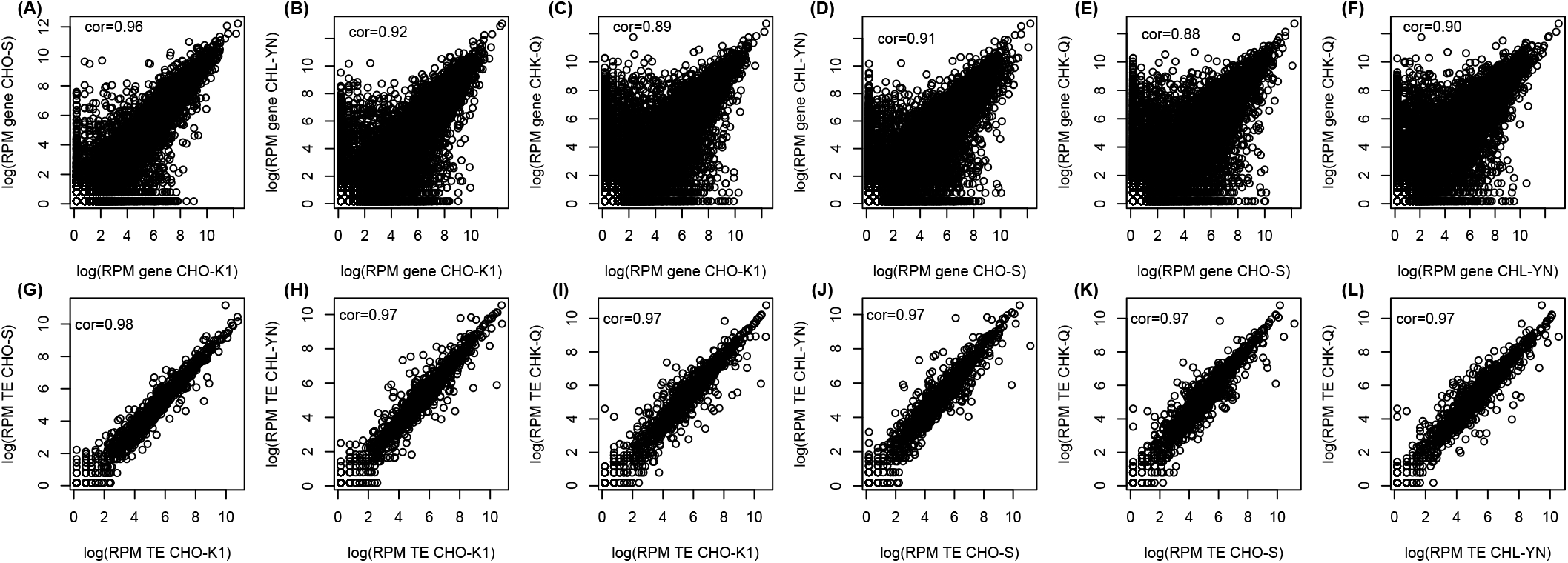
Comparisons of gene and trans elements expression between cells.

We observed numerous reference genome-dependent viral detections during our examinations. Using the older genome (CriGri-1.0) as a reference, we detected sequences such as Homo sapiens trans-activated by hepatitis C virus core protein 2 (TAHCCP2) mRNA, complete cds (AY039043.1); Macaca fascicularis adenovirus E1B 55-kDa protein-associated protein 5 mRNA, partial cds (AY650289.1); Homo sapiens retrotransposon MSTP055, complete sequence (AY972044.1); Pollicipes pollicipes strain Pol_p_N_1 histone H3 (H3) gene, partial cds (KF484309.1); Zaliv Terpenia virus strain LEIV-21C segment L, complete sequence (KF892040.1); Tete virus strain SaAn 3518 glycoprotein precursor, gene, complete cds (KM972720.1); and additional viral sequences. When using the old genomes (CriGri-1.0 and PICRH-1.0), additional viral sequences were detected, including Mammarenavirus lunaense viral cRNA, segment L, complete sequence, strain: SLW-1 (AB972431.1); Mus musculus uncharacterized long terminal repeat, complete sequence; and valyl-tRNA synthetase (G7a) gene, complete cds (AF087141.1).

We also detected cell line specific viral sequences. For example, the Kirsten murine sarcoma virus p21 v-kis protein gene (J02228.1), the Hepatitis C virus clone 110069_1_pUC_T1cl2 polyprotein gene, partial cds (KP666487.1), the Pittosporum cryptic virus-1 RdRp gene for RNA dependent RNA polymerase, segment RNA1, isolate Pit-MAIB, (LN680393.2), the Murine leukemia virus isolate Cz524, complete genome (KU324804.1), and additional viral sequences were contained only in the sequences obtained from the CHO-K1 cells. Similarly, the Wufeng Crocidura attenuata orthonairovirus 1 isolate WFS_SheQu segment M, complete sequence (OM030322.1), the Cowpox virus strain HumGri07/1, complete genome (KC813511.1) and other viral sequences were detected exclusively in sequences from the CHO-S cells. Additionally, the Mouse genomic murine leukemia virus (MuLV) related sequence (LTR-gag) (X01616.1) was detected in sequences obtained from the CHO-K1 and CHO-S cells when analyzed using the CHL-YN, CHK-Q and PICRH-1.0 genomes.

Two Chinese hamster endogenous viruses, CHERV-1b (MN527960.1) and CHERV-2g (MN527961.1) were detected in the CHO-K1, CHO-S and CHL-YN cells. However, the number of viral reads obtained from the CHL-YN cells was extremely low. We hypothesized that these reads might result from cross-contamination in the sequencing process. To test this hypothesis, we sequenced the same CHL-YN cell RNA sample using a single-use NGS platform (Nanopore). This analysis generated 6,353,244 reads and 2,589,382,340 bases and the two endogenous viruses were not detected.

### Host Cell Protein studies

The process of proteome analyses in this study using liquid chromatography-mass spectrometry (LC-MS/MS) were conducted in three steps. First, we obtained complete peptide sequences from each referential genome. Next, we developed an ion library database using the protein sequences and data from cell supernatant samples analyzed by LC-MS/MS in data-dependent acquisition (DDA) mode. Finally, we quantified individual proteins using LC-MS/MS with the ion library database in data-independent acquisition (DIA) mode. In the first step, we considered several options: using publicly available peptide sequences, obtaining peptide sequences through homology-based methods, or deriving peptide sequences via mapping-based gene prediction using RNA-seq data.

To evaluate the dependency of proteome analyses on reference genomes, we measured culture supernatant samples using various reference peptide sequences. The results showed minimal differences in measurements across different reference peptide sequences (p = 0.9292, regression analysis). Additionally, we attempted to use a previously reported ion library (Sim et al., 2020). However, in our computational environment—a general computing system provided by Sciex (Xeon Gold 6134 CPU, 3.2 GHz dual CPU, and 64 GB memory)—the library was too large to process effectively.

To validate our database assembly method for reducing false negatives in detecting high-risk HCPs, we compared two approaches. In general proteome analyses, duplicate peptide fragments in a peptide database are automatically removed or masked in LC-MS/MS systems, which can increase false negatives for HCPs. To address this, protein samples from the supernatant of CHL-YN cells were analyzed using two methods: the normal method (standard peptide sequences) and a new method in which peptide sequences of high-risk HCPs were unified. As a result, 62 out of 86 high-risk HCPs were detected using the normal method, while 74 out of 86 were detected using the new method (p = 0.4911, Fisher’s exact test). This approach reduced false negatives by 16% (Figure 2).

**Figure 2.**
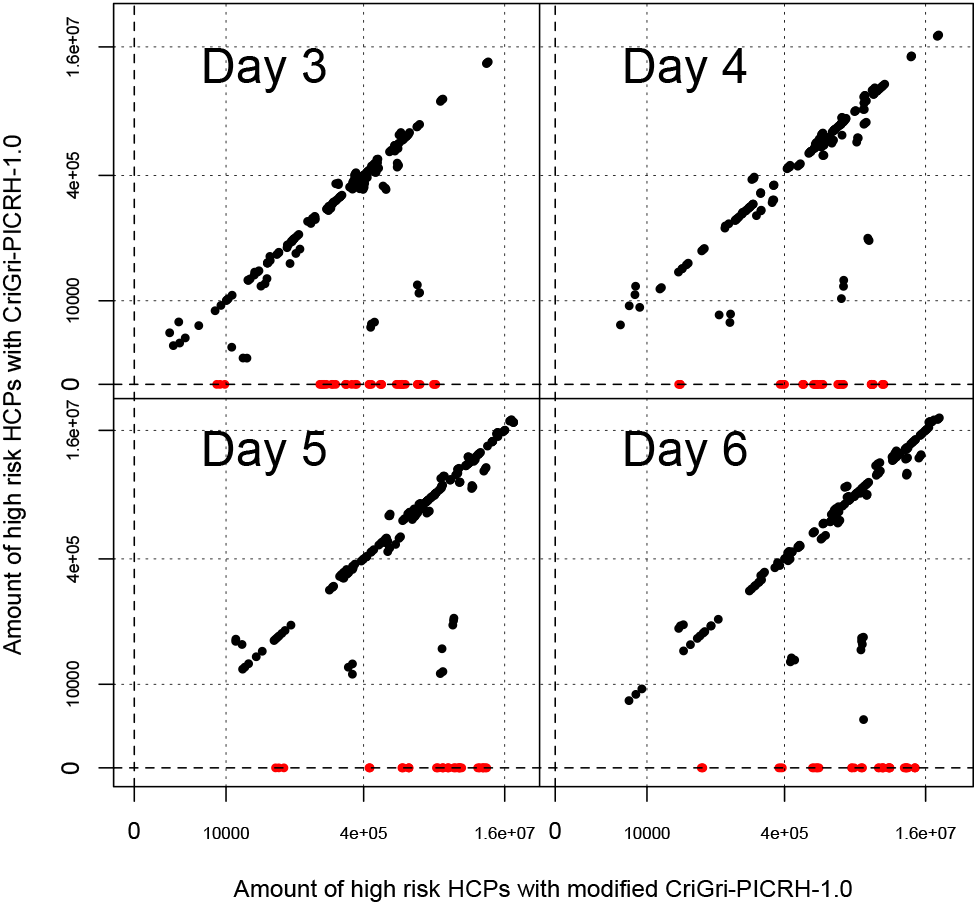
Comparison of SWATH-MS measurements using normal and modified databases. Both the vertical and horizontal axes represent the relative abundance of each protein in the samples quantified using the SWATH-MS/MS method. The vertical axis shows measurements obtained using a standard database, while the horizontal axis shows measurements obtained using a modified database. In the modified database, peptides in high risk host cell proteins were unified (CriGri-PICRH-1.0.). CHL-YN cells were cultured in flasks. The proteins undetectable in normal databases were plotted in red color.

Additionally, we measured the proteomes of the supernatant of CHL-YN cells cultured in flasks on days 3, 4, 5, and 6 (n = 3 per day). In total, 4,242 proteins were identified. The information entropy of the proteome measurements was high on days 3, 4, and 5 (10.18 to 10.22, n = 9) and decreased significantly on day 6 (9.39, 9.39, and 9.42).

### Product Heterogeneity studies

In a previous study, non-synonymous variations were identified in the recombinant humanized antibody sequence expressed in CHO cells using scRNA-seq. However, the selection of an appropriate reference genome remains unclear. In this study, we conducted a homology search between general humanized antibody sequences (OP963016.1 and OP962985.1) and Chinese hamster-derived cellular genomes. No homologous sequences were identified.

### Cellular Heterogeneity studies

In previous studies, genome expression of single Chinese hamster-derived cells was analyzed using dimension reduction techniques. In this study, we performed scRNA-seq on the C1 platform of four Chinese hamster-derived cell lines and measured genome expression through mapping and counting using six reference genomes. The annotations of newly assembled genomes were based on ab initio annotation using RNA-seq data. We also performed dimensionality reduction analyses of genome expression data. As a result, all databases functioned effectively as referential databases for numericalizing scRNA-seq data. There was little difference between two dimensionally plotted PC1 and PC2 components obtained from primary component analyses (PCA) (Figure 3). We also compared three scRNA-seq platforms (C1, 10x chromium and Rapsody) using CHL-YN cells and six genomes. Across all scRNA-seq platforms and referential genomes, we obtained similar results (Supplemental Figure 3, 4, and 5).

**Figure 3.**
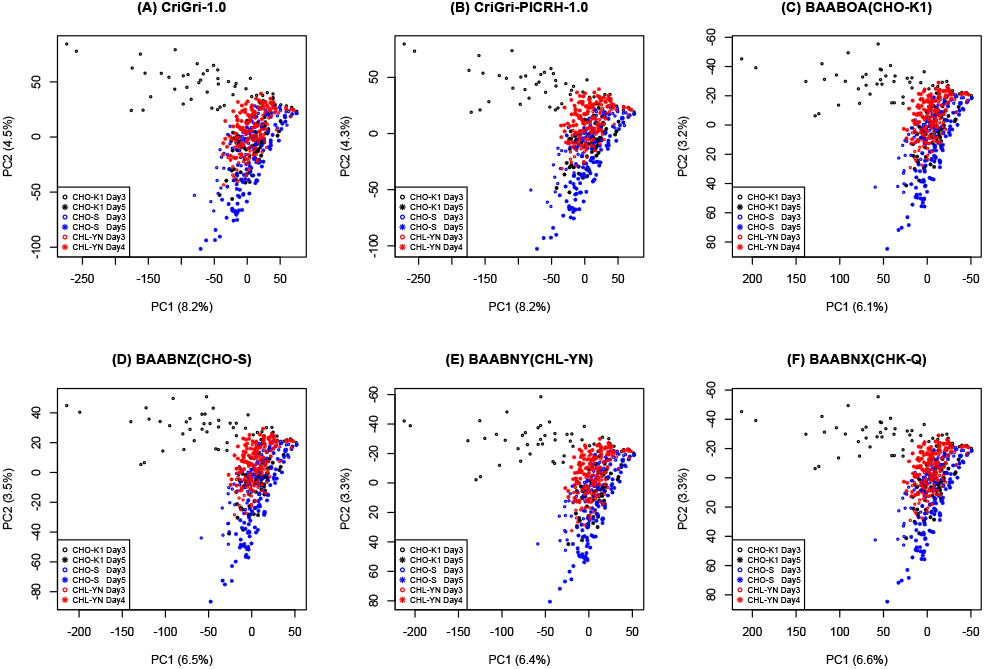
Cellular heterogeneity analyzed using several referential genomes.

**Figure 4.**
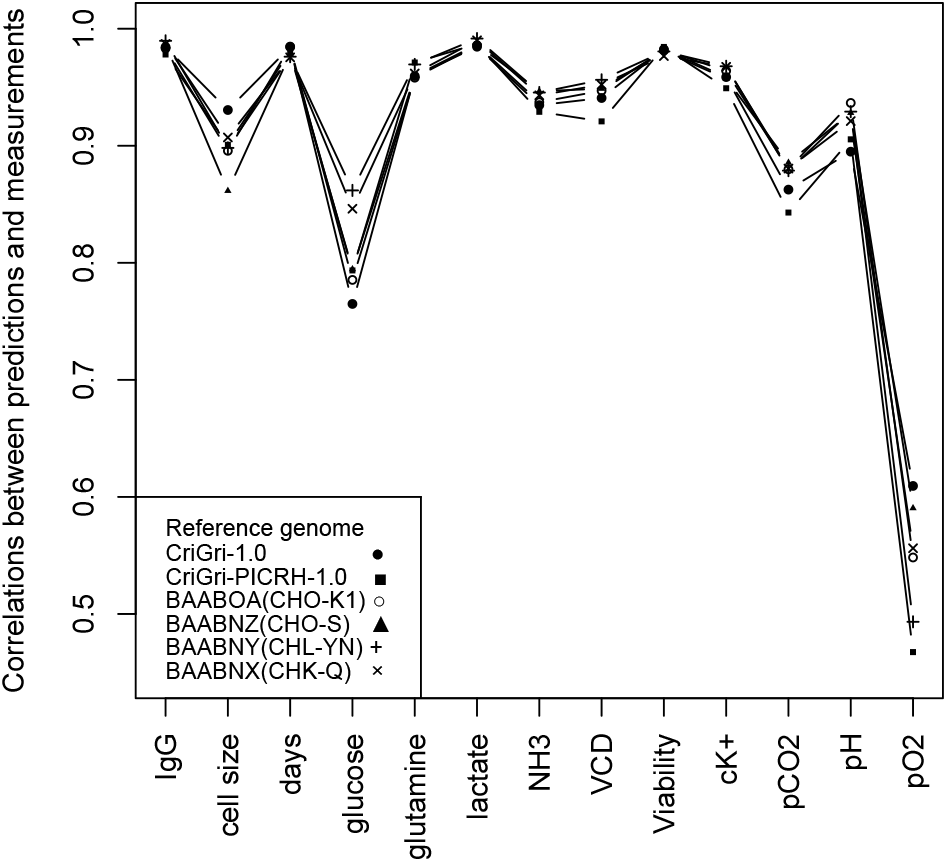
Correlations between predicted parameters and measured parameters.

**Figure 5.**
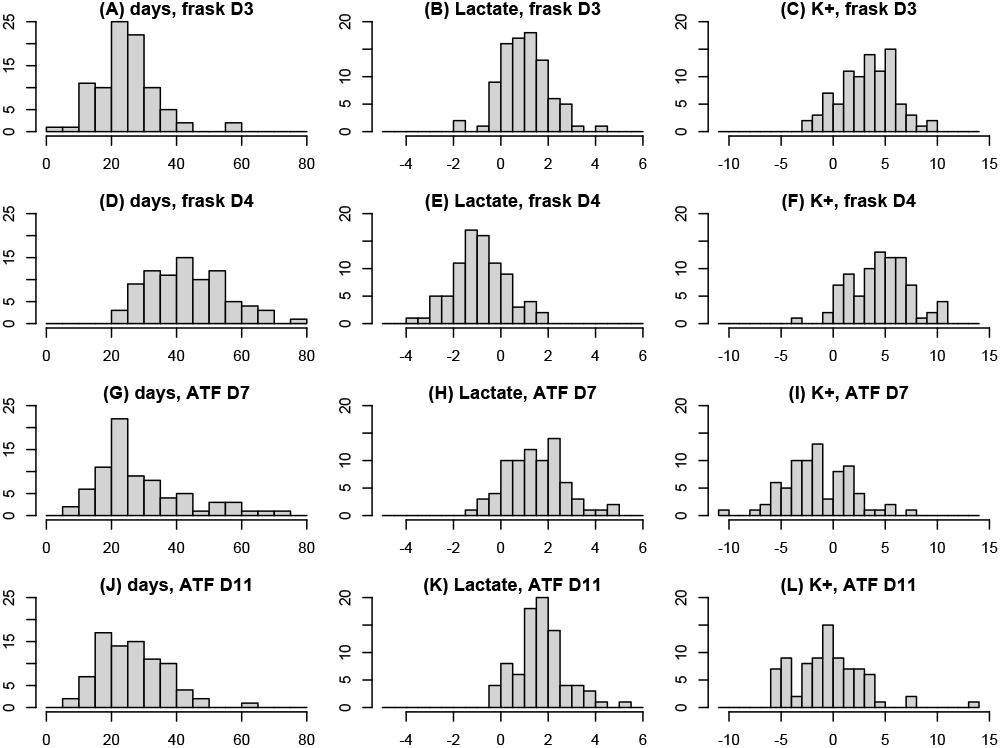
Histograms of predicted micro environmental parameters using scRNA-seq data. The histograms show the culture parameters of individual cells in each sample, predicted from single-cell RNA-seq data.The vertical axis represents the number of cells, while the horizontal axis represents the culture duration in days, lactate concentration (g/L), and potassium ion concentration (mM).Figure 4. Correlations between predicted parameters and measured parameters.

In our previous study, variants in the mitochondrial genome were developed as clonal markers of cells. In this study, we also identified variants in the mitochondrial genome. However, obtaining variants using the high-cell-number scRNA-seq platforms (10x chromium and Rapsody) was challenging due to low mitochondrial coverage. These platforms capture a larger number of cells compared to others. If we had limited the number of captured cells, we could have analyzed variants in the mitochondrial genome more effectively. **Process Reproducibility studies:** To evaluate the similarity of various culture processes, dimensionality reduction analysis can be both meaningful and straightforward. We performed principal component analysis (PCA) on bulk RNA-seq data from CHL-YN cells cultured under different conditions using six reference genomes. Little difference was observed in the results when analyzed with various reference genomes. Additionally, we applied machine learning to correlate culture condition data (e.g., IgG concentration, cellular average diameter (um), culture days, glucose (g/L), glutamine (mM/L), lactate (g/L), NH_3_(mM/L), viable cell density (cells/mL), cellular viability (%), cK+ (mM/L), pCO_2_ (mmHg), pH, and pO_2_) and genome expression data quantified using multiple genomes. The prediction accuracy was independent of the referential genomes (p = 0.999, regression analysis) (Figure 4). We also performed micro environmental prediction from scRNA-seq data using a model constructed with bulk transcriptome data and bulk environmental data (Figure 5).

**Supplemental Figure 3.**
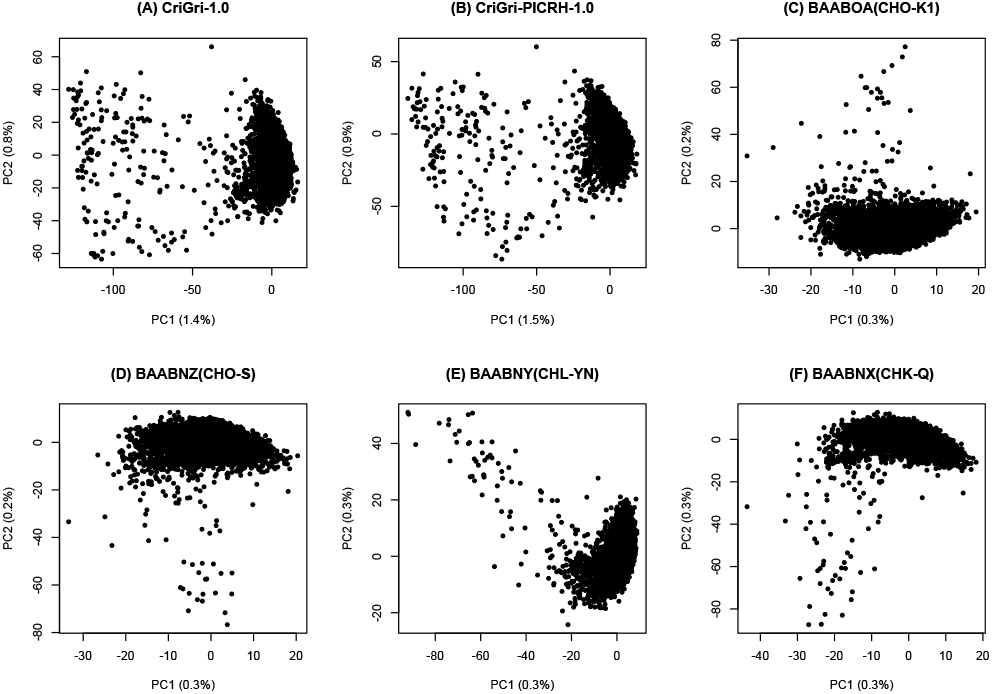
Cellular heterogeneity observed using different referential genomes in PCA using the Rhapsody platform.

**Supplemental Figure 4.**
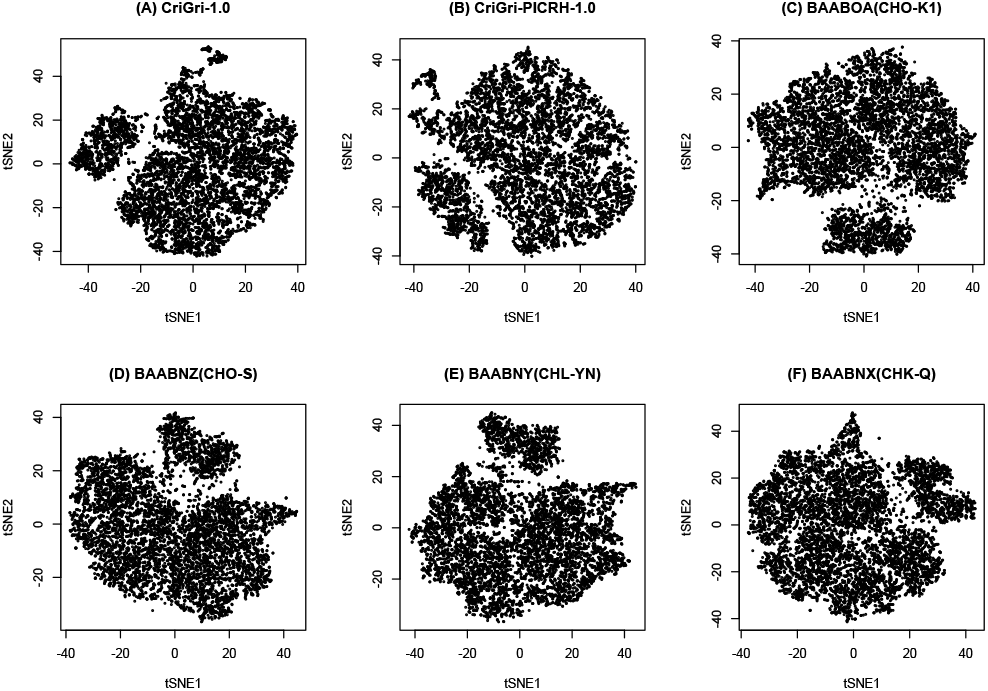
Cellular heterogeneity observed using different referential genomes in t-SNE using the Chromium platform.

**Supplemental Figure 5.**
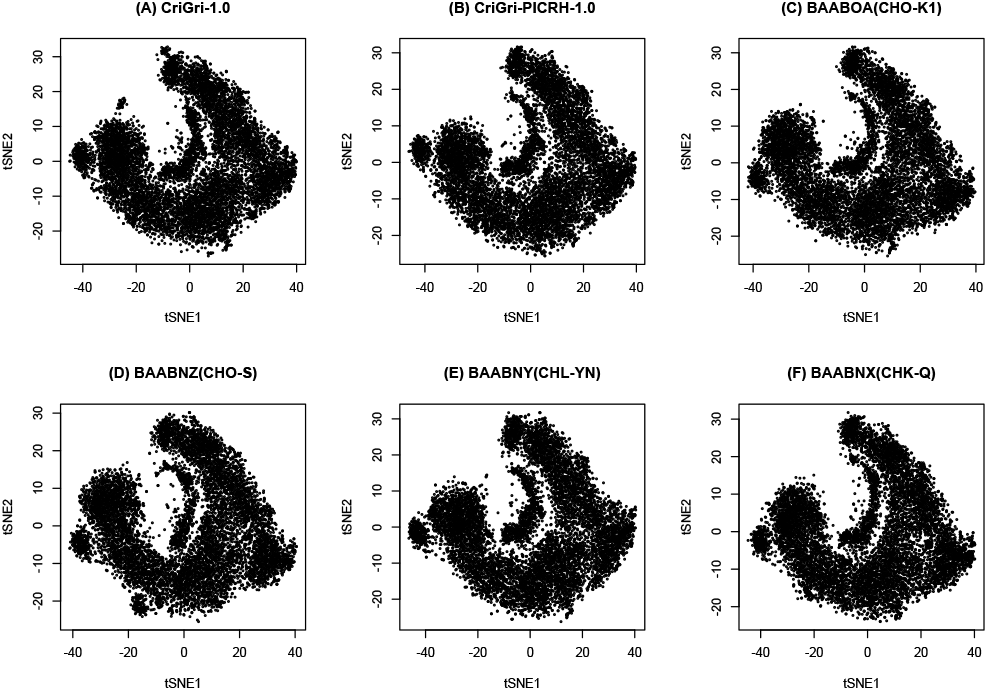
Cellular heterogeneity observed using different referential genomes in t-SNE using the Rhapsody platform.

### Ultrastructures observations

We observed over 10 cells of each cell line. We could find morphology of virus-like particles in the CHO-K1 and may in the CHO-S cells and not in the others. We also observed culture time dependent mitochondrial swelling during continuous culture (Figure 6).

**Figure 6.**
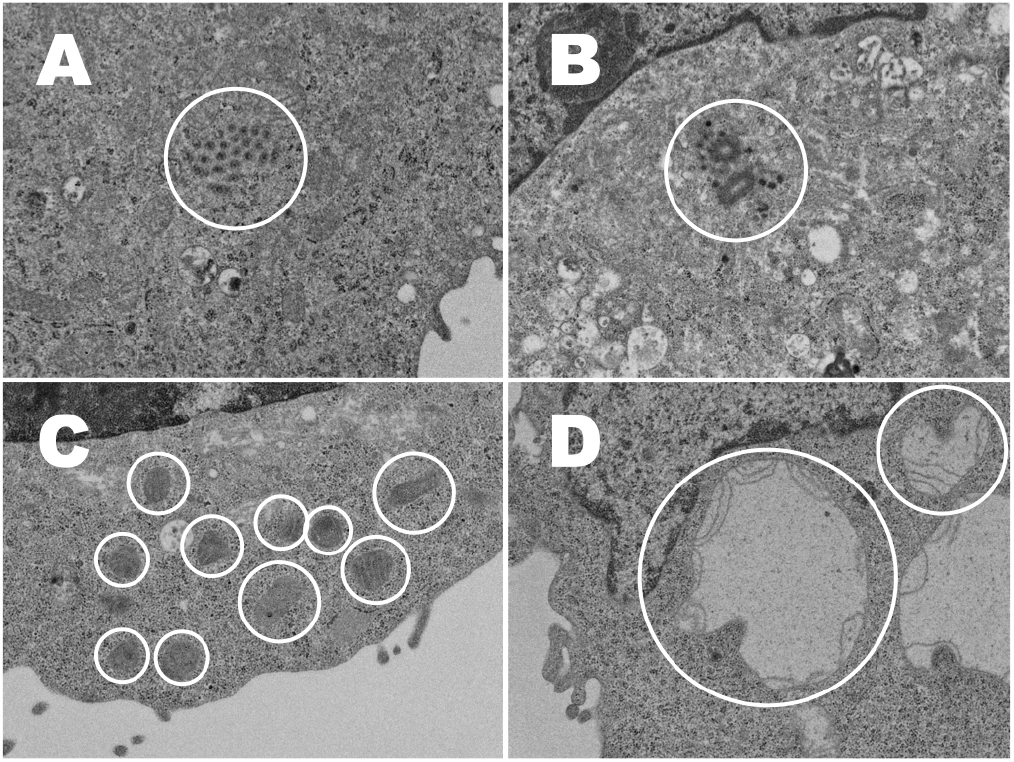
Ultrastructure of Chinese hamster cells. The figure size is 4.23 μm (height), 5.65 µm (wide). (A) a CHO-K1 cell. White circle encloses virus particles. (B) a CHO-S cell. Objects enclosed by white circle may be virus particles. (C) a CHL-YN cell. White circles enclose normal mitochondria. (D) a CHL-YN cell in a continuous culture. White circles enclose swelling mitochondria.

## IV. Discussion

Referential genomes remain essential for genomics projects, despite the development of several transcriptome analysis methods that are independent of referential genomes (Ogata & Hosaka, 2022; Yi et al., 2023). In this study, we found that any reference genomes and annotations can be used for HCP controls, product heterogeneity studies, cellular heterogeneity studies, and process reproducibility studies. Reference genomes are the fundamental source for various other reference databases, including transcriptome databases, protein databases, and ion library databases for proteomics using LC-MS/MS. In most proteomics projects, the goal is to identify differences between samples, such as those from healthy individuals and pathological cases. Consequently, general proteomics workflows are sufficient for detecting differences, but not for measuring specific proteins of interest. If the goal is to quantify a specific protein, a non-proteomics approach might be more suitable. However, in HCP safety studies for biological therapeutics, the premise is different. High-risk HCPs are already known, and the goal is to ensure that these high-risk HCPs are not overlooked in the samples. The general proteomics workflow excludes peptides shared between multiple proteins, as they do not contribute to identifying differences between samples. To meet our specific needs, we modified the proteomics workflow to focus on peptides that are part of high-risk HCPs, even if they are also part of other proteins. In this study, we masked and protected these peptides from exclusion. This approach was effective, as the signals from peptides potentially related to high-risk HCPs were enhanced. In viral safety studies, the newly assembled genomes provide the advantage of reducing the total number of unmapped orphan sequences.

We also found that viral detections are influenced by the donor cell lines and the reference genomes used. Some of these detections may result from viral contamination in the cell lines. Proving the complete absence of a virus in any cell line is impossible. To address the challenge of viral contamination in the cells, which are the source of the genome, we developed a new method. This method involves using two or more genomes. In this approach, we calculate the probability of using a virus-contaminated cell line-derived genome as v, and the number of genomes used in the process of removing host cell-derived sequences as n. The probability of avoiding a false negative in viral safety testing (f) is determined by the following equation.

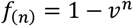

If the *v* was 0.1, then *n* = 1 is not acceptable (*f* = 0.9), whereas *n* = 2 (*f* = 0.99) is acceptable. To decrease the risk of false positive and false negative, we can use two and more genomes to annotate sequences from samples and compare the results from analyses using different genomes. If one genome contained a virus, the other genome is likely to be free from that virus. This probability-based approach to proving safety is also applied in demonstrating cellular monoclonality in the production of biological therapeutic modalities.

There are genomic differences between cell lines, including variations in genome sizes, epigenomes, and gene expression. Chromatin structure may be particularly important for bioengineering, as the nuclear incorporation of plasmid DNA occurs only during mitosis (Haraguchi et al., 2022). The newly assembled genome sizes varied, despite the k-mer distribution-based estimated genome sizes being between 2.6 and 2.7 Gb for all cell lines. The genome sizes of the two older cell lines (CHO-K1 and CHO-S) were larger than those of the newer lines (CHL-YN and CHK-Q). A genome assembly obtained from a partially immortalized cell line reflects the diversity of the cell population. The assembled genome size from a cell line may differ from the genome size of individual cells that make up the cell line. We believe this explains why the genome sizes of the older cell lines are larger, as these cell lines may exhibit greater genome diversity compared to the donor individual and/or some of the newer cell lines. Through the animal and vegetable kingdoms, nature has scattered the seeds of life abroad with the most profuse and liberal hand while nature has been comparatively sparing in the room and the nourishment necessary to rear them (Malthus, 1798). It is unclear how the natural selection processes described in Darwinism operate in in vitro environments. When observing changes in cultured cells, we must consider the effect of selection. Adequate culture media and appropriate subculturing partially free the cultured cells from selective pressures; subculturing is more of a random process than a selective one. In a future study, we may measure DAPI and/or PI intensities of the cells using FACS.

In the process reproducibility studies, we performed microenvironmental prediction from scRNA-seq data using a model constructed with bulk transcriptome and environmental data. Since we have no way to directly measure the actual microenvironmental data, these represent the most redundant microenvironmental parameters. Using this method, we can measure various microenvironments, as cells can function as micro-living sensors, similar to stingrays (Funano et al., 2020) utilizing soft sensor systems (Krehenwinkel et al., 2022). Cells in continuous cultures resemble cells in the log phase in flasks. Lactate concentration decreases in environments with uncontrolled glucose levels. Differences in potassium ion concentration are dependent on the culture media. These models are also applicable for validating sensors attached to jars and/or tanks. Therefore, measuring RNA-seq data from cells used in production is beneficial. RNA-seq data can be particularly useful when incidents occur in the process, helping to detect sensor drift, as well as viral and microbial infections. At a minimum, DNA/RNA samples should be collected and stored in freezers for commercial benefits.

## V. Conclusion

To use any referential genome is not injustice (αδικ*í*α, Ungerechtigkeit, Heidegger, 1997) in genomics-based safety and quality evaluation study of biologics, except in the case of viral safety studies. In viral safety studies, the choice of referential genome is crucial, as the number of undefined reads depends on the selected reference genome. Using two or more parallel reference genomes reduces the risk of false negative results, both theoretically and experimentally, in viral safety studies. In host cell protein (HCP) studies, this approach can also reduce the risk of false negative results for high-risk HCPs by using a peptide database in which peptides from high-risk HCPs are unified. In genomics-based safety and quality evaluation studies of biologics, database chemistry is more significant than the choice of referential genome itself (Table 1).

**Table 1.** Summary of this study.

| Object | Viral Safety | Host Cell Protein Control | Products Heterogeneity | Cellular Heterogeneity | Process Reproducibility |
| --- | --- | --- | --- | --- | --- |
| Instrument | NGS | LC-MS/MS | NGS | NGS | NGS |
| Dependence to the Quality of Genome | High | Low | Low | Low | Low |
| Progress in This Study | Use Two and More Independent Genomes | Unify the High Risk HCPs in Protein Database | - | - | Micro Environmental Prediction using scRNA-seq |

## VI. Acknowledgement

This work was funded partly by the Ministry of Economy, Trade, and Industry (METI) of Japan, and the Japan Agency for Medical Research and Development (AMED) for “Developing key technology for discovering and manufacturing pharmaceuticals used for next-generation treatments and diagnoses” (JP21ae0121014, JP21ae0121021).

## References

Baker, M. (2012). De novo genome assembly: what every biologist should know. Nature Methods, 9(4), 333–337.

Barone, P. W., Wiebe, M. E., Leung, J. C., Hussein, I. T. M., Keumurian, F. J., Bouressa, J., Brussel, A., Chen, D., Chong, M., Dehghani, H., Gerentes, L., Gilbert, J., Gold, D., Kiss, R., Kreil, T. R., Labatut, R., Li, Y., Müllberg, J., Mallet, L., … Springs, S. L. (2020). Viral contamination in biologic manufacture and implications for emerging therapies. Nature Biotechnology, 38(5), 563–572.

Birzele, F., Schaub, J., Rust, W., Clemens, C., Baum, P., Kaufmann, H., Weith, A., Schulz, T. W., & Hildebrandt, T. (2010). Into the unknown: expression profiling without genome sequence information in CHO by next generation sequencing. Nucleic Acids Research, 38(12), 3999–4010.

Bolin, S. R., Ridpath, J. F., Black, J., Macy, M., & Roblin, R. (1994). Survey of cell lines in the American Type Culture Collection for bovine viral diarrhea virus. Journal of Virological Methods, 48(2-3), 211–221.

Borsi, G., Motheramgari, K., Dhiman, H., Baumann, M., Sinkala, E., Sauerland, M., Riba, J., & Borth, N. (2023). Single-cell RNA sequencing reveals homogeneous transcriptome patterns and low variance in a suspension CHO-K1 and an adherent HEK293FT cell line in culture conditions. Journal of Biotechnology, 364, 13–22.

Campbell, M. S., Holt, C., Moore, B., & Yandell, M. (2014). Genome Annotation and Curation Using MAKER and MAKER-P. Current Protocols in Bioinformatics / Editoral Board, Andreas D. Baxevanis … [et Al.], 48, 4.11.1–4.11.39.

Cartwright, J. F., Anderson, K., Longworth, J., Lobb, P., & James, D. C. (2018). Highly sensitive detection of mutations in CHO cell recombinant DNA using multi-parallel single molecule real-time DNA sequencing. Biotechnology and Bioengineering, 115(6), 1485–1498.

Carvalho, S. B., Profit, L., Krishnan, S., Gomes, R. A., Alexandre, B. M., Clavier, S., Hoffman, M., Brower, K., & Gomes-Alves, P. (2024). SWATH-MS as a strategy for CHO host cell protein identification and quantification supporting the characterization of mAb purification platforms. Journal of Biotechnology, 384, 1–11.

Dobin, A., Davis, C. A., Schlesinger, F., Drenkow, J., Zaleski, C., Jha, S., Batut, P., Chaisson, M., & Gingeras, T. R. (2013). STAR: ultrafast universal RNA-seq aligner. Bioinformatics, 29(1), 15–21.

Funano, S.-I., Tanaka, N., Amaya, S., Hamano, A., Sasakura, T., & Tanaka, Y. (2020). Movement tracing and analysis of benthic sting ray (Dasyatis akajei) and electric ray (Narke japonica) toward seabed exploration. SN Applied Sciences, 2(12), 2142.

Graf, T., Seisenberger, C., Wiedmann, M., Wohlrab, S., & Anderka, O. (2021). Best practices on critical reagent characterization, qualification, and life cycle management for HCP immunoassays. Biotechnology and Bioengineering, 118(10), 3633–3639.

Haraguchi, T., Koujin, T., Shindo, T., Bilir, Ş., Osakada, H., Nishimura, K., Hirano, Y., Asakawa, H., Mori, C., Kobayashi, S., Okada, Y., Chikashige, Y., Fukagawa, T., Shibata, S., & Hiraoka, Y. (2022). Transfected plasmid DNA is incorporated into the nucleus via nuclear envelope reformation at telophase. Communications Biology, 5(1), 78.

Heidegger, M. (1997). De spreuk van anaximander. Leuven University Press.

ICH. (1998a). Guidance on Viral Safety Evaluation of Biotechnology Products Derived from Cell Lines of Human or Animal Origin, Q5A. International Conference on Harmonisation of Technical Requirements for Registration of Pharmaceuticals for Human Use.

ICH. (1998b). [引用] Guidance on viral safety evaluation of biotechnology products derived from cell lines of human or animal origin—FDA notice Q5A. Federal Register.

Jones, M., Palackal, N., Wang, F., Gaza-Bulseco, G., Hurkmans, K., Zhao, Y., Chitikila, C., Clavier, S., Liu, S., Menesale, E., Schonenbach, N. S., Sharma, S., Valax, P., Waerner, T., Zhang, L., & Connolly, T. (2021). “High-risk” host cell proteins (HCPs): A multi-company collaborative view. Biotechnology and Bioengineering, 118(8), 2870–2885.

Kawabe, Y., & Kamihira, M. (2022). Novel cell lines derived from Chinese hamster kidney tissue. PloS One, 17(3), e0266061.

Kim, D., Paggi, J. M., Park, C., Bennett, C., & Salzberg, S. L. (2019). Graph-based genome alignment and genotyping with HISAT2 and HISAT-genotype. Nature Biotechnology, 37(8), 907–915.

Kostic, A. D., Ojesina, A. I., Pedamallu, C. S., Jung, J., Verhaak, R. G. W., Getz, G., & Meyerson, M. (2011). PathSeq: software to identify or discover microbes by deep sequencing of human tissue. Nature Biotechnology, 29(5), 393–396.

Kovaka, S., Zimin, A. V., Pertea, G. M., Razaghi, R., Salzberg, S. L., & Pertea, M. (2019). Transcriptome assembly from long-read RNA-seq alignments with StringTie2. Genome Biology, 20(1), 278.

Krehenwinkel, H., Weber, S., Künzel, S., & Kennedy, S. R. (2022). The bug in a teacup-monitoring arthropod-plant associations with environmental DNA from dried plant material. Biology Letters, 18(6), 20220091.

Malthus, T. (1798). An Essay on the Principle of Population: The Original 1798 Edition. Suzeteo Enterprises.

Manchanda, N., Portwood, J. L., 2nd, Woodhouse, M. R., Seetharam, A. S., Lawrence-Dill, C. J., Andorf, C. M., & Hufford, M. B. (2020). GenomeQC: a quality assessment tool for genome assemblies and gene structure annotations. BMC Genomics, 21(1), 193.

Matsuda, T. (2021). Importance of experimental information (metadata) for archived sequence data: case of specific gene bias due to lag time between sample harvest and RNA protection in RNA sequencing. PeerJ, 9, e11875.

Matsuda, T., Suzuki, H., & Ogata, N. (2020). Phylogenetic analyses of the severe acute respiratory syndrome coronavirus 2 reflected the several routes of introduction to Taiwan, the United States, and Japan. In arXiv [q-bio.GN]. arXiv. 10.48550/ARXIV.2002.08802

Merten, O.-W. (2002). Virus contaminations of cell cultures - A biotechnological view. Cytotechnology, 39(2), 91–116.

Mikheenko, A., Saveliev, V., Hirsch, P., & Gurevich, A. (2023). WebQUAST: online evaluation of genome assemblies. Nucleic Acids Research, 51(W1), W601–W606.

Nanopore sequencing. (2024). Nature Biotechnology, 42(5), 698.

Nieuwenhuis, T. O., Yang, S. Y., Verma, R. X., Pillalamarri, V., Arking, D. E., Rosenberg, A. Z., McCall, M. N., & Halushka, M. K. (2020). Consistent RNA sequencing contamination in GTEx and other data sets. Nature Communications, 11(1), 1933.

Ogata, N. (2021). Whole-Genome Sequence of the Trypoxylus dichotomus Japanese rhinoceros beetle. microPublication Biology, 2021. 10.17912/micropub.biology.000487

Ogata, N., & Hosaka, A. (2022). Cellular liberality is measurable as Lempel-Ziv complexity of fastq files. 2022 IEEE 22nd International Conference on Bioinformatics and Bioengineering (BIBE), 321–326.

Ogata, N., & Iwabuchi, K. (2017). Relevant principal factors affecting the reproducibility of insect primary culture. In Vitro Cellular & Developmental Biology. Animal, 53(6), 532–537.

Ogata, N., Nishimura, A., Matsuda, T., Kubota, M., & Omasa, T. (2021). Single-cell transcriptome analyses reveal heterogeneity in suspension cultures and clonal markers of CHO-K1 cells. Biotechnology and Bioengineering, 118(2), 944–951.

Ogata, N., Shina, A., Komiya, T., Iizuka, Y., Matsuse, K., Imaizumi, F., Suwa, T., & Teramoto, A. (2018). An Electrical Impedance Biosensor Array for Tracking Moving Cells. 2018 IEEE SENSORS, 1–4.

Oh, Y. H., Mendola, K. M., Choe, L. H., Min, L., Lavoie, A. R., Sripada, S. A., Williams, T. I., Lee, K. H., Yigzaw, Y., Seay, A., Bill, J., Li, X., Roush, D. J., Cramer, S. M., Menegatti, S., & Lenhoff, A. M. (2024). Identification and characterization of CHO host-cell proteins in monoclonal antibody bioprocessing. Biotechnology and Bioengineering, 121(1), 291–305.

Patro, R., Duggal, G., Love, M. I., Irizarry, R. A., & Kingsford, C. (2017). Salmon provides fast and bias-aware quantification of transcript expression. Nature Methods, 14(4), 417–419.

Rupp, O., MacDonald, M. L., Li, S., Dhiman, H., Polson, S., Griep, S., Heffner, K., Hernandez, I., Brinkrolf, K., Jadhav, V., Samoudi, M., Hao, H., Kingham, B., Goesmann, A., Betenbaugh, M. J., Lewis, N. E., Borth, N., & Lee, K. H. (2018). A reference genome of the Chinese hamster based on a hybrid assembly strategy. Biotechnology and Bioengineering, 115(8), 2087–2100.

Shumate, A., & Salzberg, S. L. (2021). Liftoff: accurate mapping of gene annotations. Bioinformatics, 37(12), 1639–1643.

Sim, K. H., Liu, LC.-Y., Tan, H. T., Tan, K., Ng, D., Zhang, W., Yang, Y., Tate, S. & Bi, X. (2020). A comprehensive CHO SWATH-MS spectral library for robust quantitative profiling of 10,000 proteins. Scientific data, 7(1), 263.

Tanemura, H., Kitamura, R., Yamada, Y., Hoshino, M., Kakihara, H., & Nonaka, K. (2023). Comprehensive modeling of cell culture profile using Raman spectroscopy and machine learning. Scientific Reports, 13(1), 21805.

Tarailo-Graovac, M., & Chen, N. (2009). Using RepeatMasker to identify repetitive elements in genomic sequences. Current Protocols in Bioinformatics / Editoral Board, Andreas D. Baxevanis … [et Al.], Chapter 4, 4.10.1–4.10.14.

R Core Team. (2024). R: A language and environment for statistical computing. R Foundation for Statistical Computing, Vienna, Austria. URL http://www.R-Project.Org.

Tzani, I., Herrmann, N., Carillo, S., Spargo, C. A., Hagan, R., Barron, N., Bones, J., Shannon Dillmore, W., & Clarke, C. (2021). Tracing production instability in a clonally derived CHO cell line using single-cell transcriptomics. Biotechnology and Bioengineering, 118(5), 2016–2030.

Uphoff, C. C., Pommerenke, C., Denkmann, S. A., & Drexler, H. G. (2019). Screening human cell lines for viral infections applying RNA-Seq data analysis. PloS One, 14(1), e0210404.

Wang, P., & Wang, F. (2023). A proposed metric set for evaluation of genome assembly quality. Trends in Genetics: TIG, 39(3), 175–186.

Xu, X., Nagarajan, H., Lewis, N. E., Pan, S., Cai, Z., Liu, X., Chen, W., Xie, M., Wang, W., Hammond, S., Andersen, M. R., Neff, N., Passarelli, B., Koh, W., Fan, H. C., Wang, J., Gui, Y., Lee, K. H., Betenbaugh, M. J., … Wang, J. (2011). The genomic sequence of the Chinese hamster ovary (CHO)-K1 cell line. Nature Biotechnology, 29(8), 735–741.

Yamano-Adachi, N., Arishima, R., Puriwat, S., & Omasa, T. (2020). Establishment of fast-growing serum-free immortalised cells from Chinese hamster lung tissues for biopharmaceutical production. Scientific Reports, 10(1), 17612.

Yan, X., Dong, X., Wan, Y., Gao, D., Chen, Z., Zhang, Y., Zheng, Z., Chen, K., Jiao, J., Sun, Y., He, Z., Nie, L., Fan, X., Wang, H., & Qu, H. (2024). Development of an in-line Raman analytical method for commercial-scale CHO cell culture process monitoring: Influence of measurement channels and batch number on model performance. Biotechnology Journal, 19(1), e2300395.

Yi, H., Lin, Y., Chang, Q., & Jin, W. (2023). A fast and globally optimal solution for RNA-seq quantification. Briefings in Bioinformatics, 24(5). 10.1093/bib/bbad298

Yoshida, Y., Koutsovoulos, G., Laetsch, D. R., Stevens, L., Kumar, S., Horikawa, D. D., Ishino, K., Komine, S., Kunieda, T., Tomita, M., Blaxter, M., & Arakawa, K. (2017). Comparative genomics of the tardigrades Hypsibius dujardini and Ramazzottius varieornatus. PLoS Biology, 15(7), e2002266.

